# TALAIA: A 3D visual dictionary for protein structures

**DOI:** 10.1101/2022.11.28.517961

**Authors:** Mercè Alemany-Chavarria, Jaime Rodríguez-Guerra, Jean-Didier Maréchal

## Abstract

**Summary:** Graphical analysis of the molecular structure of proteins can be very complex. Full atom representations retain most of the geometric information but are generally crowded and key structural patterns can be difficult to catch. Non-full atom representations could be more instructive on physicochemical aspects but be insufficiently detailed regarding shapes (e.g., entity beans-like models in coarse grain approaches) or individualistic properties of amino acids (e.g., representation of superficial electrostatic properties). TALAIA aims at providing another layer of structural representations. It is a visual dictionary where each amino acid is represented by a unique object with differentiated shapes and colors. It makes it easier to spot important molecular information including patches of amino acids or key interactions between side chains. Most of the conventions used in TALAIA are common in chemistry and biochemistry so that experimentalists and modelers can rapidly grasp the meaning of any TALAIA depictions.

**Motivation:** The aim of the work is to offer a visual grammar that combines simplistic representations of amino acids while retaining their general geometry and physicochemical properties.

**Results:** We propose a tool that renders protein structures and encodes both structure and physicochemical aspects as a simple visual grammar. The approach is fast, highly informative, and simple, allowing the identification of possible interactions, hydrophobic patches, and other characteristic structural features at a simple glance.

**Availability:** https://github.com/insilichem/talaia

## Introduction

Molecular sciences are intrinsically related to our understanding of the three-dimensional nature of molecules. Graphical models, like those provided by computer software, serve this purpose by allowing us to perceive, describe, analyze, demonstrate, and communicate molecular knowledge including morphological, and dynamic properties, as well as reactivity. These representations stand on a series of conventions and simplifications that relate to forms (e.g., spheres for atoms, cylinders for bonds, etc.), colors (yellow for sulfur, red for oxygen, etc.), or textures (e.g., dot lines for weak interactions or depth of a bond for simple, double, or triple).

For large systems like proteins, full atom models are frequently difficult to interpret, while non-full atom representations can hide key detailed information. For example, ribbon representation of the backbone may obscure interactions of the oxygen and/or nitrogen atoms of the peptide chain. Physicochemical maps, like electrostatic ones, could be extremely informative but miss the atomic details of the systems. A frequent solution consists of allying both full-atom models with physicochemical maps. However, those hybrid representations could rapidly lead to cluttered modeling and blurred visualization. Graphical representations that combine morphological (meaning maintaining most of the shape of the amino acids), and physicochemical information are mostly absent from structural biology to date. The unique exception may be the Symbol Nomenclature for Glycans (SNFG) (Varki et al. 2015), a visual code proposed for saccharides.

Here, we propose TALAIA as a visual dictionary for proteins. TALAIA provides a visual representation of amino-acid side chains that merges structural and some physicochemical information and could serve in the inspection of protein structures, evaluation of interactions, or understanding of the driving forces of dynamical processes (e.g., a folding process). TALAIA can be applied to static structures as well as ensembles like trajectories obtained from molecular dynamics simulations.

## Availability and Implementation

The first version of TALAIA is available as a UCSF Chimera (Pettersen et al. 2004) extension and can be downloaded from the GitHub repository at https://github.com/insilichem/talaia (See repository to install). Expansion towards other platforms is underway.

In brief, the program reads the residue identity and the atomic coordinates from the input file of the protein structure to display the corresponding figures on top of Chimera’s representation. They can overlap easily with the representation of the secondary structure as well as with atomic representations (Figure 1). The transparency of the figures can be set manually.

**Figure 1.**
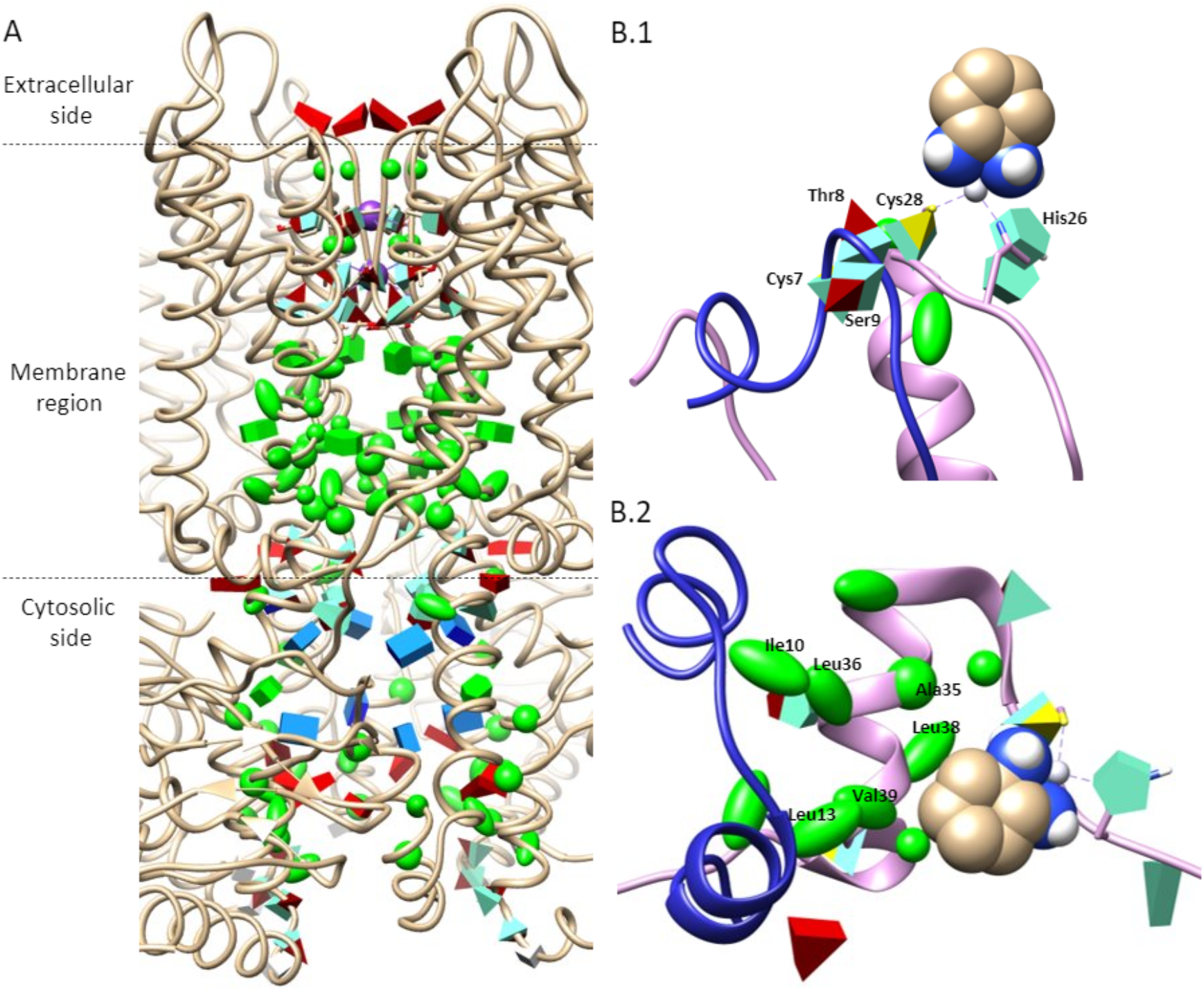
Panel A: cAMP-bound potassium channel MloK1 (PDB: 6eo1). Residues inside the channel are represented with TALAIA. Panel B: Snapshots from the molecular dynamics simulation of the insulin-oxaliplatin complex with TALAIA’s representation for residues in 8Å range from drug, showing initial conformation (B.1) and final conformation (B.2).

## 3. Description and Illustrative cases

### 3.1 Dictionary overview

TALAIA classifies amino acids into 5 families: hydrophobic, neutral polar residues, positively and negatively charged residues, and proline (Table 1). Most of TALAIA’s elements are based on conventions that could easily be accepted by molecular scientists. Regarding colors, hydrophobic residues are depicted in green, positively charged groups in dark blue, and negatively charged ones in red. Regarding the forms, they tend to reproduce the most critical aspects of the geometry of the amino acids’ side chains. Details are provided below.

**Table 1:**
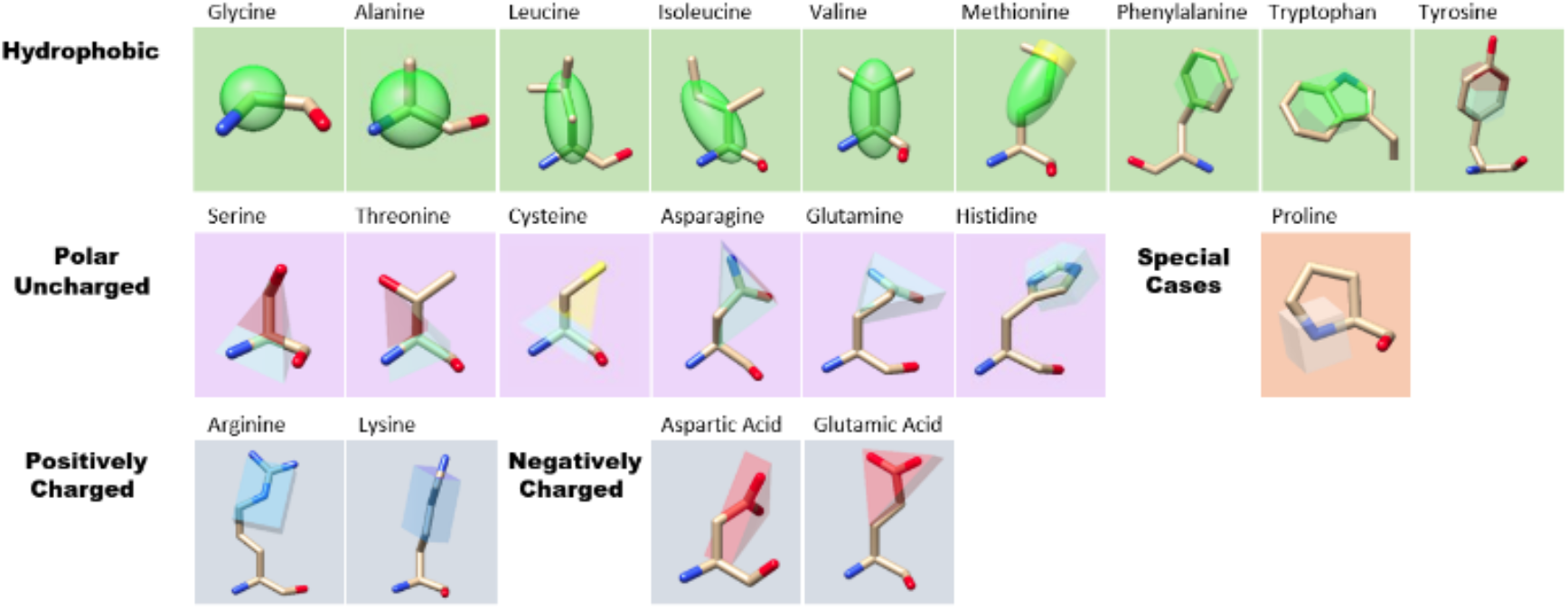
Visual dictionary classification

Aliphatic groups are presented in spheric (Glycine, Alanine), ellipsoid (Valine, Leucine, Isoleucine, Methionine), or hexagonal (Phenylalanine, Tryptophan) shapes. The elongation of the ellipsoid and hexagons are functions of their dimensions and take place along the main axis of the amino acids. Methionine, though, has a yellow ring around the ellipsoid’s end where the sulfur atom is located.

Neutral polar side chains, Serine, Threonine, and Cysteine are represented by a pyramid to indicate the directionality of the polarity. The vertex points to the polar group and the center is located on the Cα. The tip of the pyramid is yellow for Cys (representing the thiol group) and red for Ser and Thr (representing the hydroxyl group). Glutamine and Asparagine are represented by triangular prisms. The base of the prism is colored dark blue and red and located over the polar atoms of the amide group. Tyrosine is represented by a hexagon that overlaps with the residue’s ring and the hydroxyl moiety is colored in red.

Histidine is depicted by an aquamarine pentagon overlapping the residue’s ring when the residue is found in the neutral state. The color of the entire figure can change according to its protonation state, dark blue when fully protonated and red when fully deprotonated.

Positively charged residues Lysine and Arginine are represented by a blue rectangular box. The side of the cuboid where the amino group (Lys) and guanidyl group (Arg) lie are colored in a darker shade of blue. Aspartate and Glutamate side chains are depicted with the same figure as their amide forms Asparagine and Glutamine and where the carboxylic group is found is depicted in a darker shade of red.

Proline is a particular case since its structural characteristics and physicochemical properties can’t be included in the previous groups. Proline is therefore provided with a clearly distinctive code as a small grey cube centered on the Cα. This shape is chosen due to the residue’s structural role.

### 3.3 Illustrative cases

For the sake of the presentation of TALAIA to the community, we selected two cases for which a reduced number of static pictures are already informative. We selected the potassium channel of *Mesorhizobium japonicum* with PDB code 6eo1 (Kowal et al. 2018) and two snapshots from the conformational transition of the oxaliplatin once bound to insulin obtained by molecular dynamics (Sciortino et al. 2019).

Regarding the potassium channel (Figure 1.A), TALAIA allows us to immediately identify subregions: 1) a patch of negatively charged residues at the extracellular entrance of the pore, 2) small residues and neutral polar residues at the core (the selectivity filter), and 3) a hydrophobic region formed mainly by S6 and C-linker at the end of the transmembrane region. This representation allows fast recognition of a ring of positively charged arginine residues where the cyclic nucleotide-binding domain binds near the membrane interface. Below that ring, negatively charged residues are located as well marking the cytosolic end of the pore.

Regarding the insulin-oxaliplatin system, figure 1.B presents two snapshots of the conformational transition observed in a molecular dynamics simulation for one of the low-energy complexes obtained by protein-ligand dockings (Sciortino et al. 2019). This complex results from the cleavage of Cys28-Cys7 disulfide bridge so that Cys28 could coordinate the platinum ion together with His26. TALAIA depiction allows us to rapidly highlight that during the transition, the second coordination sphere interaction of the metal with Thr8 and Ser9 is lost (Fig. 1.B.1). This allows the N_Ter_ part of the helix to unfold, and finally the aromatic part of the drug can reach a hydrophobic core formed by Ile10, Leu13, Ala35, Leu36, Leu38, Val39 (Fig. 1.B.2).

We believe that TALAIA provides an interesting complementary tool for the molecular inspection of biomolecular systems by affording a visual code that preserves global geometric blueprints of amino acids that are physicochemically sound.

